# Tissue-specific Hi-C analyses of rice, foxtail millet and maize suggest non-canonical function of plant chromatin domains

**DOI:** 10.1101/567883

**Authors:** Pengfei Dong, Xiaoyu Tu, Haoxuan Li, Jianhua Zhang, Donald Grierson, Pinghua Li, Silin Zhong

**Author notes:** These authors contributed equally to this work (Pengfei Dong, Xiaoyu Tu). To whom correspondence should be address: Silin Zhong and Pinghua Li.

## Abstract

Chromatins are not randomly packaged in the nucleus and their organization plays important roles in transcription regulation. Using *in situ* Hi-C, we have compared the 3D chromatin architectures of rice mesophyll and endosperm, foxtail millet bundle sheath and mesophyll, and maize bundle sheath, mesophyll and endosperm tissues. We have also profiled their DNA methylation, open chromatin, histone modification and gene expression to investigate whether chromatin structural dynamics are associated with epigenome features changes. We found that plant global A/B compartment partitions are stable across tissues, while local A/B compartment has tissue-specific dynamic that is associated with differential gene expression. Plant domains are largely stable across tissues, while rare domain border changes are often associated with gene activation. Genes inside plant domains are not conserved across species, and lack significant co-expression behavior unlike those in mammalian cells. When comparing synteny gene pairs, we found those maize genes involved in gene island chromatin loops have shorter genomic distances in smaller genomes without gene island loops such as rice and foxtail millet, suggesting that they have conserved functions. Our study revealed that the 3D configuration of the plant chromatin is also complex and dynamic with unique features that need to be further examined.

## Introduction

How chromatin is packaged inside the nucleus has fascinated biologists for a long time. Recent advances in chromosome conformation capture technologies such as Hi-C have enabled us to examine the spatial organization of chromatin architectures (Yu and Ren 2017). At the genome-wide scale, the mammalian Hi-C contact map shows a checkerboard plaid-like pattern, suggesting that chromatins are not randomly packaged inside the interphase nucleus, and some are more likely to interact with one another (Lieberman-aiden et al. 2009). Principal component analysis can classify chromatin regions into two different compartments based on the interaction pattern. The “A” compartments are euchromatin with active transcription, open chromatin and active epigenome marks, while the “B” compartments are heterochromatin with transposable elements, repressive chromatin marks and DNA methylation (Lieberman-aiden et al. 2009). At the sub-megabase scale, mammalian chromatin could be further divided into topologically associating domains (TADs) by analyzing their local interaction pattern (Nora et al. 2012; Dixon et al. 2012). TADs are considered to be the structural-functional unit of the mammalian genome, and are highly stable and even conserved across species (Sexton and Cavalli 2015; Dixon et al. 2016).

Extensive studies have now confirmed that TADs could influence gene expression by facilitating spatial interactions of cis-regulatory elements and genes in the same TAD, while preventing them from interacting with those in other TADs (Sexton and Cavalli 2015; Dixon et al. 2016). It has been shown in many cases that deletion or duplication of TAD boundaries could lead to gene mis-regulation due to abnormal interactions of enhancer cis-regulatory element and proximal gene promoter (Lupiáñez et al. 2015; Franke et al. 2016; Hnisz et al. 2015; Narendra et al. 2016). In addition, multi-scale computational interrogation of TADs and their insulation potential has confirmed that TAD represents a functionally privileged scale of chromatin architecture, arising from their ability to partition chromatin interactions (Zhan et al. 2017). Crosslinking-free 3C approach also confirmed that the restrictions imposed by TADs are present in the native nuclei, and they are not artifacts of crosslinking (Brant et al. 2016). Finally, several recent studies have identified structural proteins that are required for the establishment of TAD boundaries, and perturbations of those factors could cause ectopic gene expression, strongly arguing for the functional roles of TADs (Zuin et al. 2014; Haarhuis et al. 2017; Nora et al. 2017; Rao et al. 2017; Schwarzer et al. 2017).

TAD-like domains have been reported in several plant species including cotton, maize, tomato, sorghum, foxtail millet and rice (Liu et al. 2017; Wang et al. 2017a; Dong et al. 2017, 2018). Whether plant domains have the same biological function as their mammalian counterparts, however, remains largely unknown. It is difficult to use conventional genetics to study them in plants, particularly without prior knowledge of factors that could affect their establishment or function, such as the mammalian CTCF and cohesin complex (Haarhuis et al. 2017; Rao et al. 2017; Nora et al. 2017). However, we do know the consequences of mammalian TAD formation, which is to regulate gene expression within the domain. For example, genes located within the same TADs have higher expression correlation than genes located in different TADs (Nora et al. 2012). Both human and mouse cell-type and tissue-specific Hi-C analysis showed that genes located in the same TAD are co-regulated (Le Dily et al. 2014; Zhan et al. 2017). In addition, TADs are also highly conserved suggesting they have important function that are under strong selection (Vietri Rudan et al. 2015; Dixon et al. 2012).

To examine whether plant chromatin structure could have similar biological functions to the mammalian ones, we have constructed tissue-specific Hi-C maps of multiple plant species, including leaf mesophyll and endosperm cells of rice, leaf mesophyll and bundle sheath tissues of foxtail millet, and mesophyll, bundle sheath and endosperm tissues of maize (Table S1). We then associated these chromatin 3D architecture data with tissue-specific epigenome and transcriptome profiles.

## Results

### Plant A/B compartment partition is stable across tissues

We first examined whether the genome-wide A/B compartments could change in different tissues. Unlike the mammalian chromosomes that could be partitioned into multiple active “A” and repressive “B” compartments by principal component analysis of the correlation of the genome-wide Hi-C matrix (Lieberman-aiden et al. 2009), plant chromosomes often adopt a simple structure with two A compartments at the chromosome tips and one continuous B compartment in the center (Fig 1) (Dong et al. 2017; Liu et al. 2017). The two A compartments correspond to the euchromatin arms, while the central B compartment is the pericentromeric heterochromatin. In some cases, like the tomato chromosomes, large gene islands inserted into the pericentromeric heterochromatin could also form A compartments and interact with the two A compartments in the euchromatin arms. After examining the Hi-C matrix of different plant tissues, we found that their chromosome A/B compartment partition pattern is highly stable across all plant tissues we examined (Fig 1, Table S6). This is different from the dynamic mammalian A/B compartment partition that is functionally related to transcription activity and often displayed tissue-specific changes (Dixon et al. 2015; Schmitt et al. 2016).

**Figure 1.**
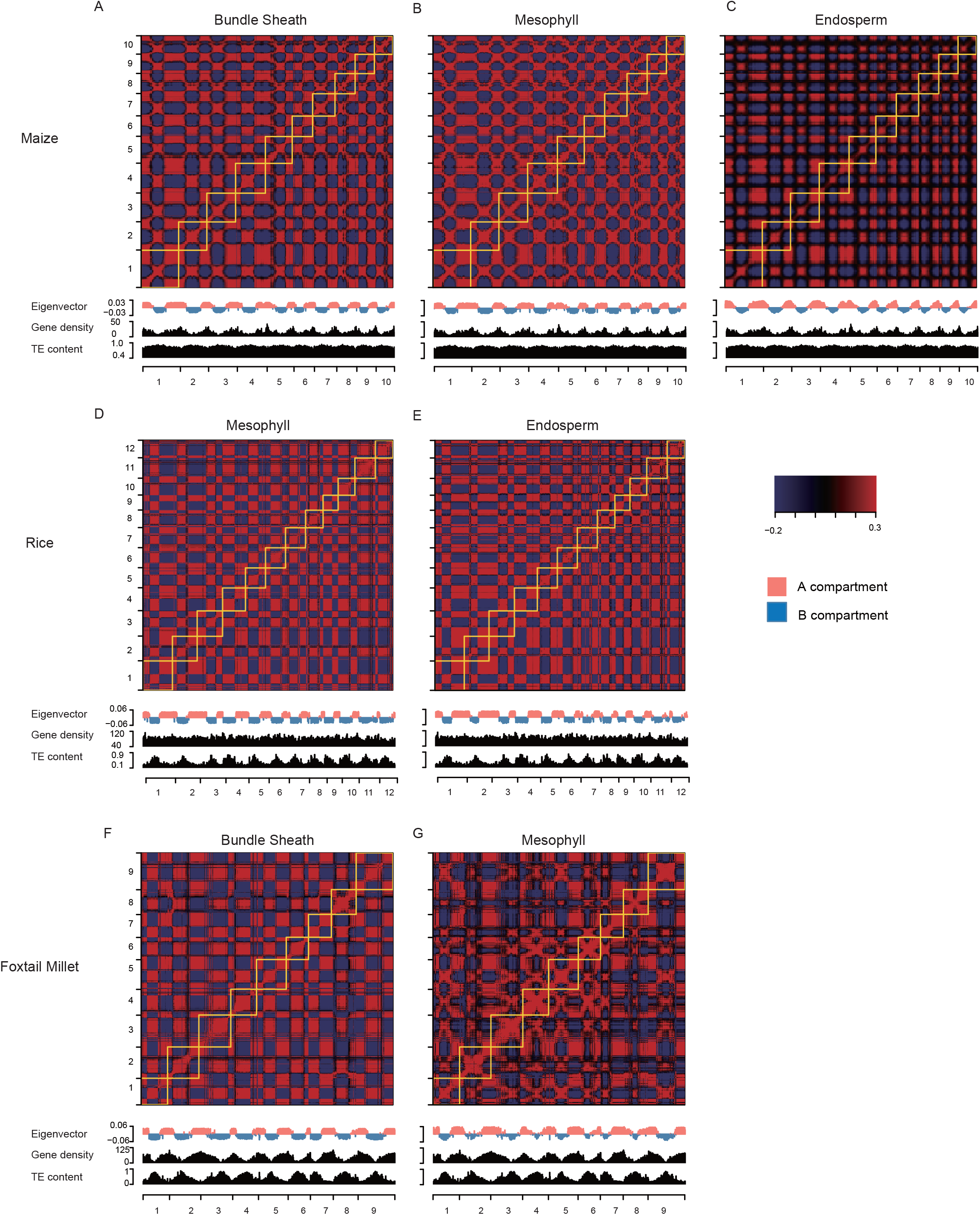
Global A/B compartment is stable across different tissues. Pearson correlation matrix illustrates the correlation between the intra- and inter-chromosomal interaction profiles. The eigenvector and genome feature tracks are shown below the matrix. a) Maize bundle sheath, mesophyll and endosperm tissues. b) Rice mesophyll and endosperm tissues. c) Foxtail millet bundle sheath and mesophyll tissues. Unlike the mammalian ones, plant A/B compartment partition is closely associated with the euchromatin and heterochromatin states of the chromosome and lack tissue-specific dynamics.

Although the plant A/B compartment partition is stable, we observed that chromatin contact intensity between B compartments is weakened in the rice and maize endosperm tissues (Fig 2). For example, in the rice endosperm tissue, the interaction between B compartments decreased 13.7% (p<2.2e-16, Wilcoxon rank sum test) when compared to those in the leaf mesophyll tissue (Fig 2C). The same pattern was found in the maize endosperm tissue when compared to both leaf bundle sheath and mesophyll tissues. The reduction is severe enough to be visible from their genome-wide Hi-C contact matrices. (Fig 2 A and B). We also compared the interaction decay exponents of all tissues, which is represented by the steepness of the slope, with which chromatin interaction frequencies decay with genomic distance. Although the Hi-C signal is the sum of all maternal and paternal chromatins in the endosperm tissues, the decreased chromatin interaction frequency is consistent with previous studies that the maternal genome in endosperm tissue is hypomethylated and their heterochromatin is decondensed (Bauer and Birchler 2006; Feng et al. 2010).

**Figure 2.**
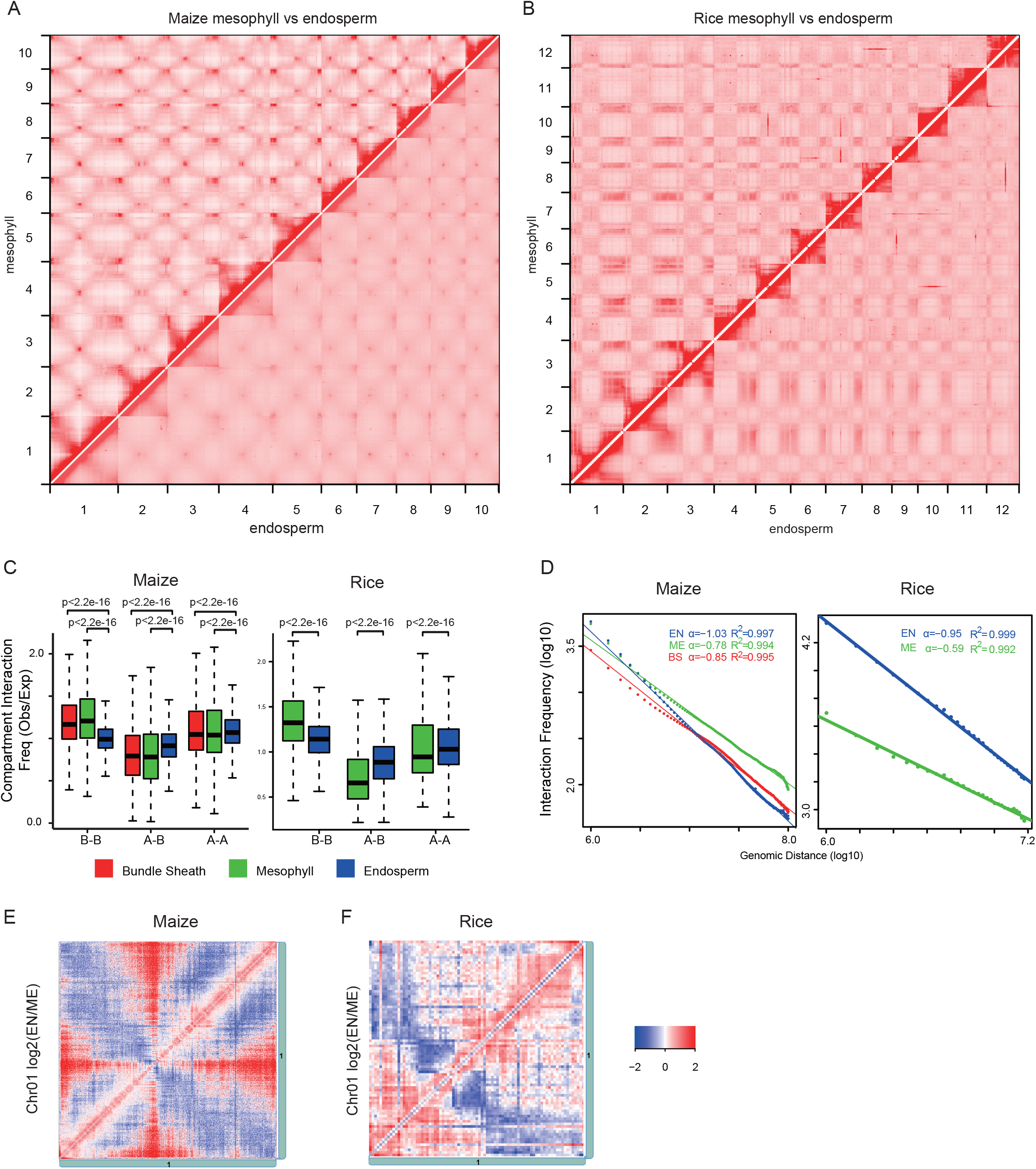
Reduction of global B compartment interaction in the endosperm. Genome-wide Hi-C contact maps of mesophyll vs endosperm tissues in (a) maize and (b) rice. c) Global A/B compartment interaction frequency (Wilcoxon rank sum test). d) Interaction decay exponents of different maize and rice tissues. e-f) Local chromosomal Hi-C matrix showing the ratio of endosperm to mesophyll contact frequency.

A unique feature of the Hi-C contact map of plants with large genome size is the intense interaction signal on the anti-diagonal lines forming the X-shaped pattern (Mascher et al. 2017; Dong et al. 2017). This indicates that their chromosomes could be organized as a U-shape, and the chromosome arms could intact with each. When we examined the maize endosperm Hi-C map at the chromosome level, we found that the interaction frequency on the anti-diagonal line is reduced and those between other regions became more intensed (Fig 2 A and E). Rice, however, lacks this X-shaped pattern (Fig 2B). The decreased contact frequency between its chromosome arms in endosperm is still visible, which is indicated by the strong blue color on in the anti-diagonal line (Fig 2F).

### Local (sub) A/B compartments dynamics associated with transcription

For large plant genomes, conventional A/B compartment partition methods are unable to separate the active gene islands from the largely inactive heterochromatin environment. We have previously shown that large plant chromosomes could be further divided into local A/B compartments, also referred to as sub-compartments (Doğan and Liu 2018; Dong et al. 2017). This partition method better reflects the local active and repressive status of the plant chromatin. Therefore, we examined whether local A/B compartment would display tissue-specific dynamics (Fig 3).

**Figure 3.**
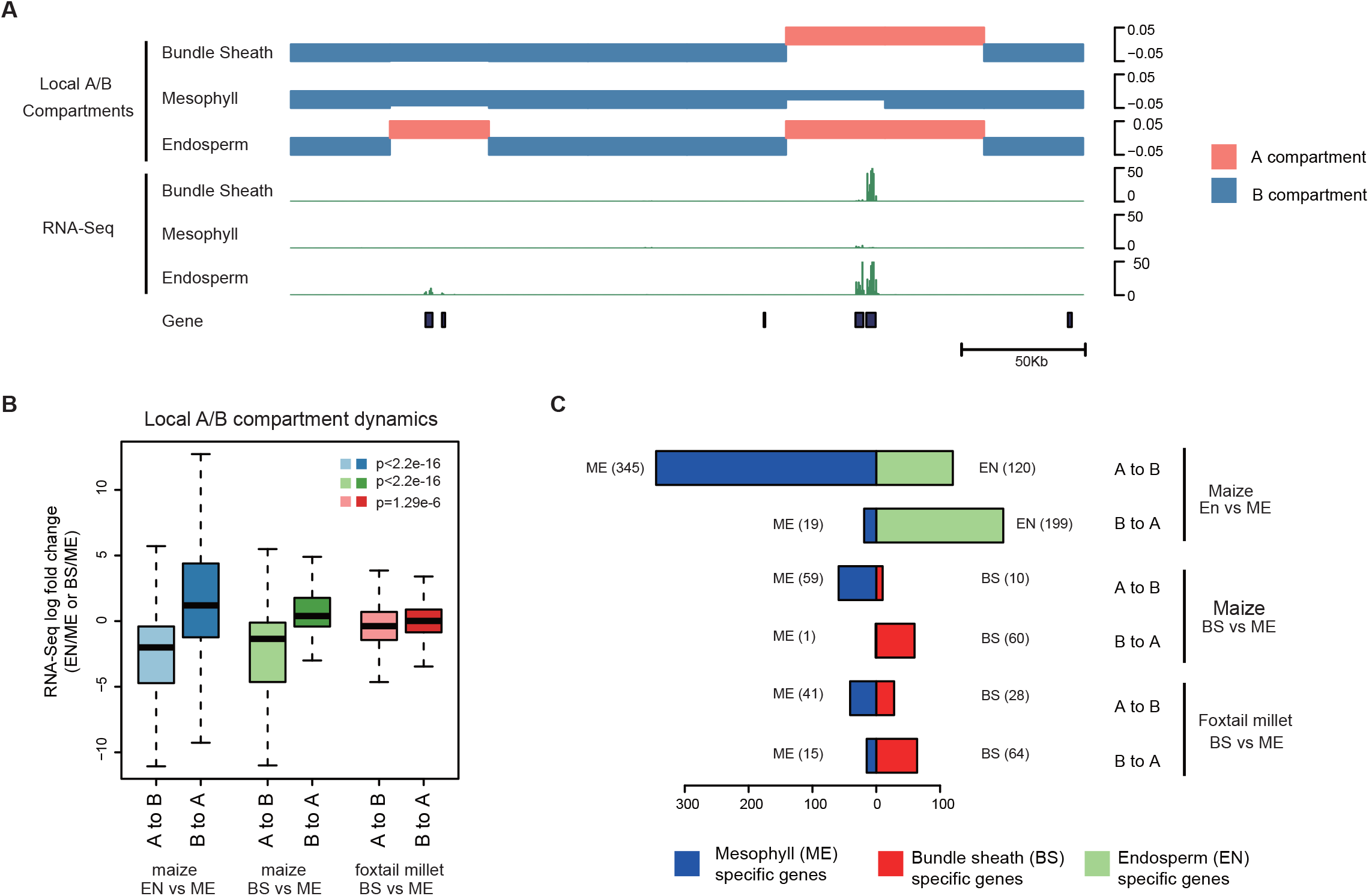
Local A/B compartment change associated with transcription. a) An example of local A/B compartment change in maize (Chromosome 2: 30.48 to 30.80 Mb). The gene expression tracks are shown below the eigenvector. b) Switches from local A to B compartment are associated with reduced RNA-Seq signal, and *vise versa* (Wilcoxon rank sum test). c) Box plot showing region associated with tissue-specific local A/B compartment changes are often associated with tissue-specific gene expression.

In total, we have identified 24.02 Mb and 176.52 Mb regions that experienced tissue-specific local A/B compartment changes in foxtail millet and maize, respectively (Table S7). We found that the inactive B to active A local compartment changes are indeed associated with transcriptional activation, while the A to B local compartment changes are associated with transcriptional repression (Fig 3B and C). Consistent with the observed reduction of global B compartment interaction in maize endosperm (Fig 2C-E), we found that its local compartment partition is also weakened (Fig S2 C). The rice endosperm chromatin can hardly be partitioned into local compartments, and this is clearly visible from the local Hi-C correlation matrix that lacks a checkerboard pattern (Fig S2 F).

To quantify and confirm this reduction of local chromatin interaction, we first divided their genomes into the same 20 and 40 kb bins that are used for local compartment analysis in rice and maize, respectively. Next, we inferred their active or repressive state using the normalized RNA-Seq read counts and TE density in each bin. We compared the interaction pattern of the most active bins (top 10% RNA-Seq signal) and the most repressive bins (top 10% TE content) (Table S8, Figure S2). We found that the average normalized interaction frequency between the repressive regions significantly dropped from 1.30 in rice mesophyll to 1.12 in endosperm (p<2.2e-16, Wilcoxon rank sum test). The same pattern was observed when we calculated the endosperm interaction frequency among regions previously identified as local B compartment in leaf tissues (Table S9, Fig S2). These results suggest that the repressive regions, no matter whether they could be partitioned into local B compartment or not, are less compact in the endosperm tissues.

### Plant domains are more stable than mammalian ones

Mammalian TADs are considered to be stable across tissues and even species (Dixon et al. 2012, 2015). For example, 65.53% (1289/1967) of TAD border regions in human embryonic stem cells have been identified as border regions in fibroblast cells (Dixon et al. 2012). Plant domains are not conserved across species. With the new tissue-specific Hi-C data, we first examined whether the plant domain boarders are conserved between tissues. In maize, we found 85.75% and 78.99% of the mesophyll domain border regions are still identified as domain border regions in bundle sheath and endosperm tissues, respectively (Fig 4). We observed high correlation of the border region insulation scores of different tissues, the local minima of which are used for domain border calling (Fig 4C).

**Figure 4.**
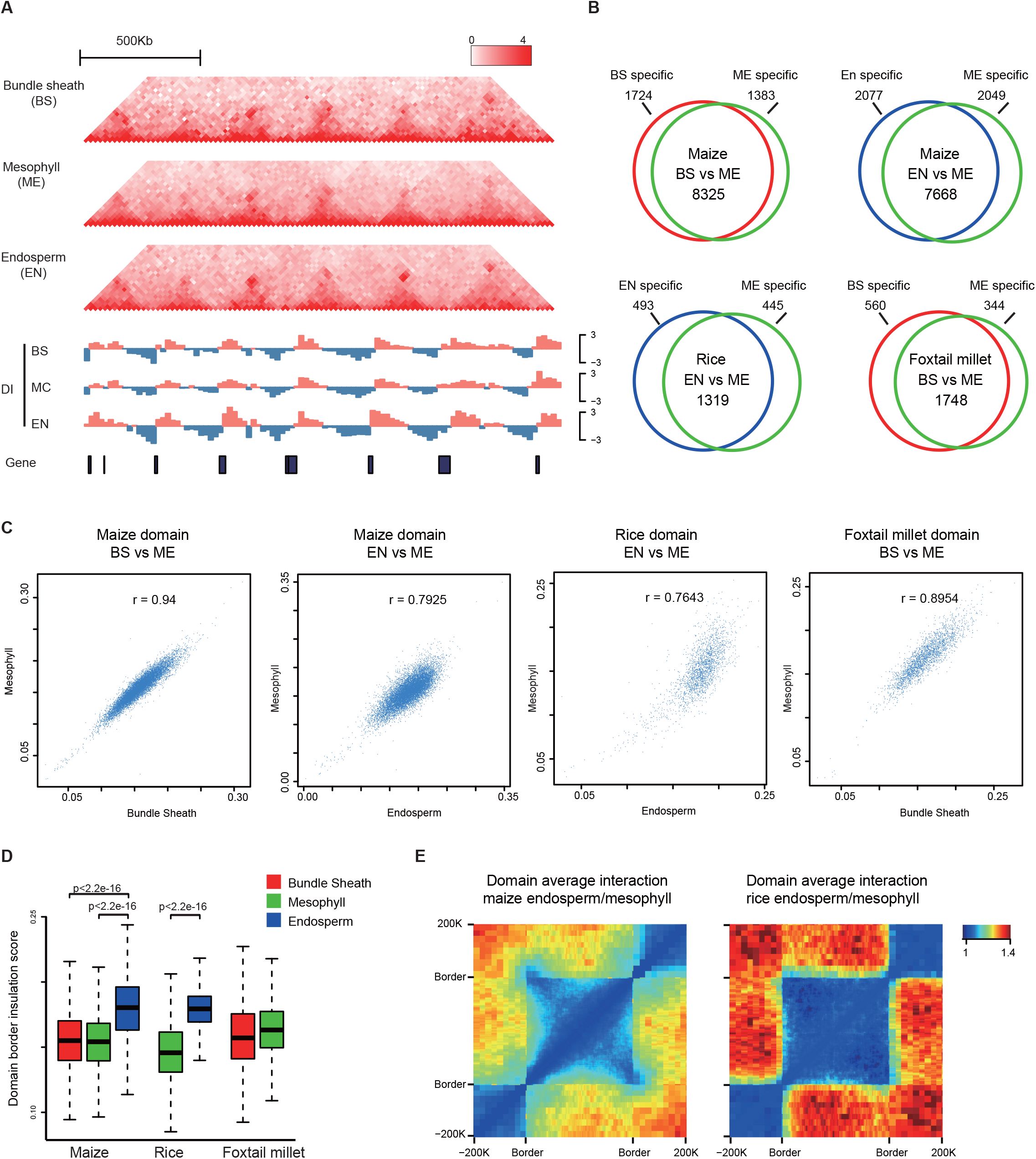
Plant domains are stable across tissues. a) Example of domain organization in maize (Chromosome 10: 20-22Mb). The directionality index (DI) is shown below the Hi-C contact map. b) Number of tissue-specific domain borders in each species. c) Pearson correlation coefficient of insulation scores in the domain border region. d) Despite domains being stable, the endosperm tissues have higher domain border insulation scores (Wilcoxon rank sum test). e) Heatmap showing the ratio of the endosperm to mesophyll average contact frequency. Both the maize and rice endosperm have lower within domain average contact frequencies, suggesting that their domains are less compact.

Next, we examined the non-conserved maize domain borders, which we defined as regions called as border in one tissue but failed to be identified as border in another tissues. We found that their insulation scores are still local minima, and the correlations co-efficiency of the interaction directionality indexes in these border regions are significantly higher than the random regions (Fig S4). The same patterns were observed in the rice and foxtail millet domain data (Fig S4). These showed that the plant non-conserved borders still have weak border features, suggesting that plant domains are very stable across tissues.

We compared the within- and cross-domain interaction intensities between different tissues. We first divided the conserved domains into 100 bins, while the regions 200 kb up and down-stream of the domain were divided into 50 bins each. We then compared the average interaction frequency for each bin from the endosperm and mesophyll Hi-C data (Fig 4 D and E). The results showed that despite the fact that domains themselves are not changed, inter-domain interaction could vary between tissues. The endosperm has higher contact frequencies across the domain borders when compared to other tissues, which indicates that endosperm domains are less compact and is consistent with the increased border insulation score and the reduced global and local B compartment interaction. It should be noted that the reduction of within domain interaction means that the interactions across the domain border became relatively higher, leading to a higher insulation score calculated for the endosperm domains (Fig 4D and 4E).

### Plant domain border changes are associated with transcription

In mouse, it has been shown that gain of a new tissue-specific TAD boundary is often associated with gene up-regulation in the border region (Bonev et al. 2017). We found that the plant domain border change is also associated with tissue-specific gene expression (Fig 5, Fig S5). For example, we have identified 4,126 tissue specific borders in the maize endosperm and mesophyll pair-wise comparison (Fig 4 B). In the 2,077 endosperm-specific borders, we found 2.04 times more endosperm up-regulated genes than down-regulated genes, while in the 2,049 mesophyll-specific borders, 1.52 times more mesophyll up-regulated genes were found than down-regulated genes (p<2.2e-16, Fisher exact test). In other maize and rice tissue pair-wise comparisons, gain of domain border is also associated with gene up-regulation (Table S10).

**Figure 5.**
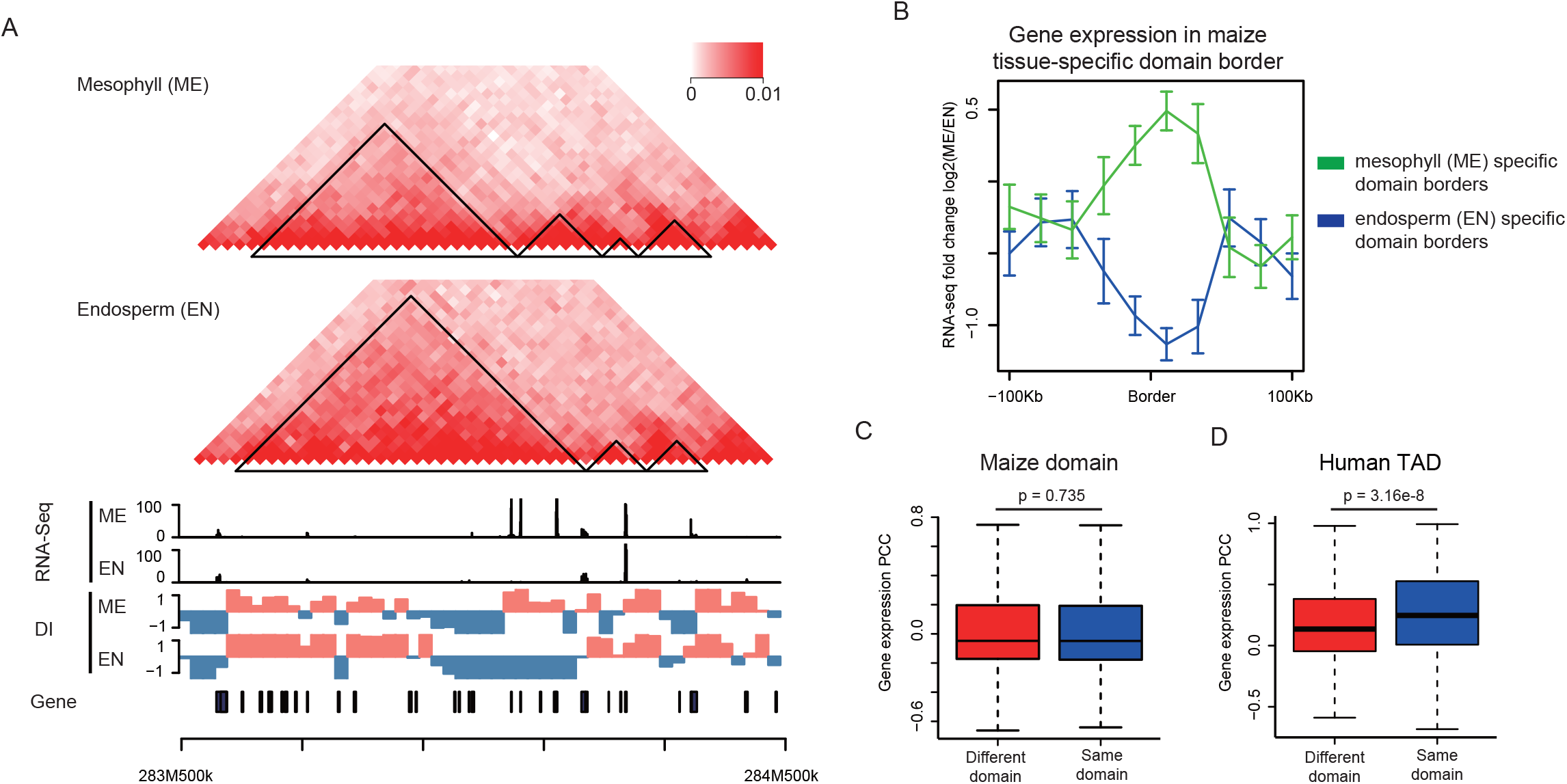
Domain border changes are associated with transcription change. a) Example of a tissue-specific domain border change in maize chromosome 1. The normalized gene expression and DI (directional index) are shown below the Hi-C matrix. b) Fold change of RNA-Seq signal in tissue-specific domain border regions. c-d) Pearson correlation coefficient of domain gene expression in maize. Human domain data was shown as a comparison (Rao et al. 2014). Only gene pairs with 100 kb physical distance were shown, the rest are shown in Fig S6.

### Plant domain genes lack significant co-expression and co-regulation patterns

Mammalian TADs can contribute to gene regulation by restricting chromatin interactions, such as those between genes and distal regulatory elements. Hence, the gene pairs within the same TADs are associated with higher gene expression correlation, when compared to pairs of the same physical distance but located in two different TADs (Nora et al. 2012). However, when we examined the expression correlation of gene pairs located within the same plant domains, they are not significantly different from the randomized control (Fig 5C, Fig S6). For example, the correlation of human gene pairs located in the same TADs is significantly higher than the control (Fig 5D), but in maize, the median of expression correlation for intra-domain pairs is not significantly different from that of the inter-domain pairs (−0.046 vs −0.0480, p=0.735, Kolmogorov-Smirnov test). The same pattern was observed when we examined other plant domain gene pairs of different physical distances. We also repeated the analysis using only the conserved maize domains, which are defined as domains identified consistently in mesophyll, bundle sheath and endosperm tissues, and the result was the same (Fig S6).

Besides having higher expression correlation in general, mammalian genes located within the same TAD tend to be co-regulated when comparing Hi-C and expression data of two tissues (Le Dily et al., 2014; Zhan et al., 2017). Therefore, we tested whether plant genes inside the same domains could still be co-regulated using the same approach. The domains were classified as up- or down-co-regulated if the number of up- or down-regulated genes within the domain is higher than 95% (p=0.05) of the randomized control domains. In the maize bundle sheath vs mesophyll comparison, 129 up-regulated domains could be identified out of 2,545 actual domains, while 114,381 up-regulated ones were found in the 2,366,066 randomized control domains (p=0.58, Fisher exact test, Table S11). A similar lack of co-regulation was observed in all tissues pair-wise comparisons we performed. This is different from the previous studies using mammalian TADs (Zhan et al. 2017).

Our results showed that genes located in plant domain are not co-regulated, nor do they tend to be associated with higher expression correlation. This could suggest that plant domains might not possess the same biological function as the mammalian TADs.

### Gene island loops are stable across tissues and distance between loop genes are under selection

The majority of the human and mouse chromatin loops detected by Hi-C are stable across tissues, and the dynamics ones are often associated with tissue-specific gene expression changes (Rao et al. 2014). In the large plant genomes such as tomato and maize, we have previously identified chromatin loop formed between highly expressed gene islands separated by condensed heterochromatins (Dong et al 2017). It should be noted that these large chromatin loops detected by Hi-C were identified at the resolution of tens of kb, and are much larger than the gene proximal promoter and distal regulatory element loops detected by ChIP-PET assay.

We first examined whether these maize gene island chromatin loops have tissue-specific dynamics like the mammalian ones (Fig 6D, Fig S7). We have used the Juicer package to identify tissue-specific loops (Durand et al. 2016). The result showed that the plant loops are also stable across different tissues (Table S12). For example, less than 5% of the loops are called as tissue-specific in the bundle sheath and mesophyll pair-wise comparison (Table S12). Similar to the results in mammalian cells, we also observed few examples of tissue-specific loops that are associated with tissue-specific gene expression patterns (Table S13).

**Figure 6.**
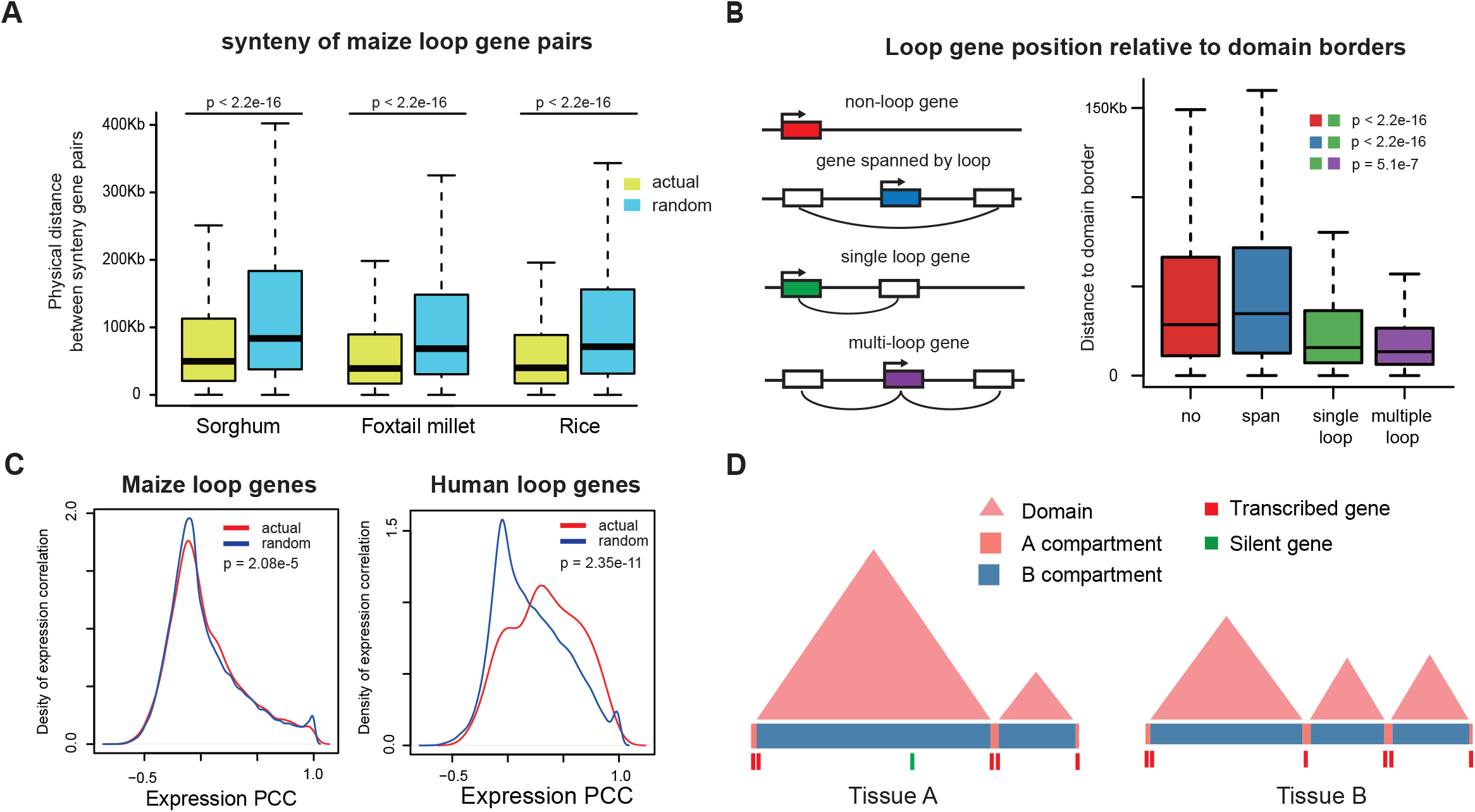
The maize chromatin loops and a model for tissue-specific chromatin 3D organization dynamics a) The physical distance of the synteny gene pairs of the maize loop genes are under selection in sorghum, rice and foxtail millet. b) Genes involved in loops are located closer to the domain borders (Wilcoxon Rank Sum test). c) Maize loop gene pairs lack a clear co-expression pattern when compared to the human ones (Kolmogorov-Smirnov test). d) Schematic representation of tissue-specific domain and local compartment change associated with gene expression.

Mammalian chromatin structures such as TAD and loops are highly conserved between cell types and even across different species, suggesting that they have important functions and are under selection (Krefting et al. 2018; Dixon et al. 2012). Hence, we examined whether the physical distance of the maize gene pairs involved in loops could be conserved and under selection in other monocot species. We first identified the synteny of the maize loop gene pairs in sorghum, rice and foxtail millets. Although we do not know whether they could form small loops that are below the detection limit of Hi-C, we can still calculate the physical distance of these synteny gene pairs. Therefore, we are able to compare the physical distance of the loop synteny pairs to the randomized control pairs, which are defined as synteny genes of random non-loop-associated maize gene pairs of the same physical distance (Fig 6A). For example, we found that the average distance of rice loop synteny gene pairs is 38,813 bp, which is 1.79 times smaller than the randomized control pairs (p<2.2e-16, Wilcoxon rank sum test). The same pattern was observed for gene-pairs of different physical distance in sorghum and foxtail millet, suggesting that these loop synteny gene pairs are indeed under selection, and might have biological functions.

We have previously analyzed that tomato and maize leaf Hi-C data, and found that their loops are enriched outside domains near the border region (Dong et al., 2017). In contrast, loops in human and mouse cells are enriched within TADs near the border regions, and they are often referred to as corner loops (Rao et al. 2014). In this study, we found the similar loop distribution pattern in the maize bundle sheath and endosperm tissues. For example, we found that maize genes involved in loop are located closer to the domain border than non-loop genes (Fig 6, Fig S8, Table S14). In addition, genes involved in multiple loops are closer to domain border than genes involved in a single loop. Genes spanned by the a chromatin loop are shorter than genes located in the two anchor regions (Figure S8).

In mammalian cells, genes involved in chromatin loops have higher expression correlation (Lan et al. 2012). However, the maize loop genes lack the strong co-expression correlation as those associated with loops in mammalian cells (Fig 6C). The same pattern was observed when we examined loop gene pairs of different physical distances in different tissues (Fig S9, Table S13).

## Discussion

It is now well recognized that 3D chromatin architecture is important for the regulation of gene expression in mammalian cells. Recent studies in different plant species have revealed that large genome plants also have complex 3D structures such as compartments, local-compartments (sub-compartments), domains and chromatin loops (Liu et al. 2017; Dong et al. 2017). However, questions remain to be addressed regarding whether plant chromatin structures have biological functions, and whether their functions are similar to those of the mammalian ones.

A/B compartments are known to have tissue-specific dynamics that are often associated with the transcription activity in mammalian cells (Dixon et al. 2015). This is not the case for plants, and we found that they are in accordance with the euchromatin and heterochromatin status, which is constant between tissue types (Fig 1). Only the local A/B compartment displayed tissue-specific dynamics that could be correlated with gene expression changes (Fig 3), suggesting that they could be equivalent to the mammalian A/B compartment. We also observed reduced interactions among endosperm B compartments and increased interaction between the chromocenter and the euchromatin arms (Fig 2). This coincides with the reported hypomethylation and decondenstation of the maternal endosperm genome (Zhang et al., 2014). Similar changes have also been reported in Hi-C analysis of the *Arabidopsis* DNA methylation mutants, which have a decondensed heterochromatin center (Feng et al. 2014). As we performed all the experiment using self-pollinated plant tissues and could not distinguish the paternal and maternal chromatins, it will be interesting to perform an allelic Hi-C analysis using a cross-pollinated endosperm or embryo tissues.

In mammalian genomes, TADs are considered to be the core structural-functional unit, which can confine the chromatin interactions between distal regulatory elements and the genes that they regulate (Sexton and Cavalli 2015; Dixon et al. 2016). Disruption of TAD border could lead to the loss-of-regulation of distal enhancers (Lupiáñez et al. 2015; Franke et al. 2016; Hnisz et al. 2015; Narendra et al. 2016). A recent study has shown that during neural differentiation, newly gained TADs are associated with gene up-regulation in the domain border region (Bonev et al. 2017). In addition, DNA recombinations are enriched in TAD borders and depleted within TADs, and genes within TADs have more conserved expression patterns (Krefting et al. 2018; Dixon et al. 2016). All these findings suggested that TADs are under selection and strongly argue for a functional role for TAD in transcriptional regulation.

If the plant domains serve similar functions as the mammalian TADs, that is to confine chromatin interaction between distal regulatory elements and proximal gene promoter, they could be under selection and conserved in related species. However, we found that plant domains are not conserved (Dong et al. 2017). In tomato and maize, large chromatin loops are formed between gene islands in gene-poor heterochromatin regions. The loop genes are positioned close to domain borders and the gene synteny pairs have significantly shorter physical distances in related species, suggesting that they are under selection (Fig 6). However, these plant loops are enriched outside the domains, suggesting that the domains might not serve as a spatial insulator that confine these large chromatin interactions. However, we need to take into account that these gene island loops detected by Hi-C are unable to reflect the interaction between gene and distal cis-regulatory elements, which are often a few kilo base pair part. The mammalian genome organization is very different from those of the plants, and their promoter to distal cis-regulatory elements loop could be detected by Hi-C.Hence, we could not rule out that plant domains might be able to confine the local chromatin loops. But these loops are unlikely to confine plant domains as the mammalian corner loops

A recent study also found that domains in rice seedling are largely stable in different growth conditions, while their domain border changes are not associated with gene expression changes (Liu et al. 2017). Here, we have compared different tissues and found that plant domains are stable across tissues, while plant genes within the same domain also lack clear co-expression pattern, unlike the mammalian model. Therefore, compare to the wealth of comparative and functional studies of chromatin 3D architectures in the animal systems, we still know too little about them in plants, not to mention their potential functions.

## Materials and Methods

### Plant materials

Rice (variety minghui 63) leaf of two-week-old seedlings and milky stage endosperm tissues were used in this study. Maize (variety B73) and foxtail millet (variety Yugu1) two weeks old seedling leaf mesophyll and bundle sheath as well as 10 days post anthesis endosperm tissues were used. To isolate mesophyll and bundle sheath cells, the lower epidermis of the leaf was peeled and submerged in enzyme solution (0.6 M sorbitol, 20 mM MES pH5.5, 1 mM CaCl_2_, 1 mM MgCl_2_, 0.1% BSA, 1% cellulose R10 and 0.2% macerozyme R10). The leaf tissues were vacuum infiltrated for 10 seconds and incubated at room temperature for 3 to 4 hours with gentle shaking. The mesophyll protoplasts were carefully flushed off the surface using a dropper, and the solution was filtered through a 60 um cell strainer. The mesophyll cells were collected by centrifugation at 100 × g for 10 min, and used for formaldehyde fixation in 1x PBS or DNA/RNA extraction. The remaining bundle sheath strands on the filter were then recovered and suspended in new enzyme solution without the cell wall digestion enzymes. The solution was vortexed vigousely to lyse the remaining undigested mesophyll cells, and the purify of the bundle sheath strands were examined under microscope. Examination of the mesophyll and bundle sheath specific marker genes in RNA-Seq and histone H3K27me3 ChIP-Seq data also confirmed the purify of the isolated cells. For example, NADP-ME was only expressed in the maize bundle sheath cells and associated with mesophyll-specific H3K27me3 marks.

Part of the isolated cells and bundle sheath strands, as well as the endosperm tissues were fixed for 15 minutes in 1xPBS with 1% formaldhyde, and their nuclei were used for Hi-C and histone ChIP-Seq with 2-3 biological replicates each (Table S1-4). For the ATAC-Seq assay, unfixed protoplast and bundle sheath strand were first lysed by adding TritonX100 to 1%, and incubate in the refrigerated rotator for 10 to 20 minutes to release the nuclei. The nuclei were then collected by centrifugation at 1000 × g for 10 min, and washed multiple times till the color changed from green to grey, which indicates the chloroplasts have been removed. The purified nuclei were then resuspended in 50 uL of 1x Tagmentation Buffer and incubated with 1 uL Tn5 for 20 min at room temperature. The reaction was then terminated by added EDTA and SDS, and the DNA was purified and PCR amplified using Illumina N70x and N50x index primers. The resulting libraries were size fractioned by agroase gel and the small fragments smaller than a mono-nucleosome (<250 bp) were purified and sequenced.

### Data processing

In this study, we have used the maize ensembl build 36, foxtail millet v2.2, and rice indica Minghui63 as reference genomes. Transposable elements were identified as previous described (Dong et al. 2017). Synteny gene pairs between maize and other grass species include foxtail millet, sorghum and rice are downloaded from CoGe (https://genomevolution.org/).

Hi-C analysis were performed as previously described (Dong et al., 2017). The reproducibility between biological replicates were calculated by HiCrep (Yang et al. 2017) (Figure S1). For rice and foxtail millet domain calling, we used 20 kb bin size. For maize domain and loop calling, we used 10 kb bin size as previously described. To confirm that the sequencing depth is sufficient under such resolution, we followed the method described in Lieberman-aiden et al. (2009), which requires the top 80% of the bins have no less than 1,000 *cis*-interaction (Figure S1).

The BS-Seq reads were aligned to reference genomes using bsmap (Xi and Li 2009). The DNA methylation ratios for each cytosine were determined by methratio.py from the bsmap suite.

For RNA-Seq analysis, sequencing reads were aligned to reference genomes using Tophat2 (Langmead et al. 2009). The read counts for each gene were obtained through HT-Seq packages (Anders et al. 2015). Normalized reads per kilobase per million mapped reads counts were calculated using the R package DESeq2(Love et al. 2014). We also used DESeq2 to identify differentially expressed genes (FDR smaller than 0.01 and log fold change greater than 2). For histone ChIP-Seq and ATAC-Seq data, the reads were mapped to reference genome using bowtie2 (Langmead and Salzberg 2012). Reads mapping quality greater than 10 were kept as unique mapped reads and PCR duplicates were removed by samtools (Li et al. 2009). Enriched regions were called using macs2 (Zhang et al. 2008) with default parameters.

### Global A/B compartment analysis

We identified the global compartments based on the first principle component of the distance normalized whole-genome interaction matrix using 500 kb bin as previously described (Dong et al. 2017). We estimated the interaction decay exponents at 500 kb resolution. We grouped loci pairs of the same genomic distance and calculated their average interaction frequency. After that, we used a linear regression model to fit the log ratio of the average interaction and the log ratio of the genomic distance. The resulting slope of the model is the desired interaction decay exponents (Fig 2D). For the larger maize genome, we examined the genomic distance between 1Mb to 100Mb, while the range of 1Mb to 15Mb is used for the smallest rice genome. Since the interaction frequency between global A and B compartment increased significantly in endosperm, we also limit the loci pair examined to be those located within a consecutive global compartment domain.

### Local A/B compartments analysis

Local compartments were identified as previously described (Dong et al. 2017). The chromosomes are first divided into blocks by constrained clustering using the normalized Hi-C contact matrix. The CscoreTool was then used to call local compartment for each block (Zheng and Zheng 2018). For maize, we used 40Kb bin, while 20 kb bin size was used for rice and foxtail millet.

Both the maize and rice endosperm tissues exhibit reduced local compartment partition, and rice endosperm chromatin could not be partitioned to local compartments. To infer and calculate their local compartment interactions, we divided their chromosome into bins, and calculate their RNA-Seq read counts and TE contents. The top 10% mRNA read counts bins as defined as active regions, and top 10% TE content bins are defined as repressive regions. We then calculated the distance normalized interaction between active-to-active (A-A), active-to-repressive (A-R), and repressive-to-repressive (R-R) regions. Only regions located in the same block were kept for the analysis. Same result could be obtained if we used local compartment positions defined in the rice leaf Hi-C dataset to calculate the interaction of endosperm.

Next, we examined regions associated with local A/B compartment change and associate them to gene expression changes. We called the tissue-specific local compartment region when it showed the same A-to-B or B-to-A change in both replicates, as well as in the merged replicates. Next, we identified the differentially expressed genes using DESeq2 and their log2 fold-change values, which were used for Wilcoxon rank sum test to compare the gene expression associated with A-to-B and B-to-A local compartment changes.

### Domain analysis

Domains were identified by hitad with 20 kb bin as previously described (Wang et al. 2017b; Dong et al. 2017). The general domain features are the same as the previously reported (Dong et al., 2017, Figure S3). To compare domain across different tissues, the domain border were considered to be conserved between tissues if the two borders are located in the same or the adjacent bins. Insulation score were calculated as previously described (Alekseyenko et al. 2015). For each 21-bins (20Kb) windows sliding along diagonal line, the insulation score S was defined as

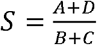

where A, B, C, and D is the sum of lower left, upper right, and lower right block matrices (10 × 10) of the square Hi-C interaction frequency matrix (21 ×21).

To compare the average inter- and intra-domain interaction in endosperm and mesophyll tissues, we isolated the Hi-C matrix of their conserved domain with 200 kb up- and down-stream regions. We divided these conserved domains into 100 equally sized bins and their 200 kb up- and down-stream regions were divided into 50 equally sized bins. We used the R approx function to perform the constant interpolation of the distance normalized interaction matrix, which first interpolated each row and then each column into 200 bins. After that, interaction matrix of the domain and its 200 kb neighboring region is normalized into the same size. We could then divide the endosperm interaction matrix by the mesophyll interaction matrix, and obtained the average ratio to be used in Fig 4 E.

To test whether genes within the same domain exhibit similar expression pattern, we obtained published RNA-Seq data (Li et al. 2010; Tobergte and Curtis 2013; Liu et al. 2013), and calculate the correlation of gene expression between gene pairs within the same domain and in different domains as previously described (Le Dily et al. 2014; Nora et al. 2012). To avoid bias caused by genomic distance of the pairs, we only examined gene-pairs of similar genomic distance from 50 kb to 100 kb. Only expressed genes (fpkm>1) in at least 3 tissues were used in this analysis. Same analysis was performed using the published human GM12878 TADs and expression data (Rao et al. 2014; Keen et al. 2015).

We examined whether genes within the same domain is up- or down-coregulated using the method previously described (Zhan et al. 2017). For the randomized control, we cyclically permutated the genome for 1000 times and calculated the up- and down-coregulated genes for the permutated domains. We then calculated the mean and standard deviation of the up-/down-regulated gene numbers for domains with the same gene number. Random domains containing more genes than the actual domains were discard to prevent bias. The Z-score is calculated as follows

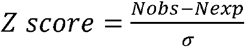

where Nobs is the observed up-/down- coregulated gene number, Nexp is the mean and σ is the standard deviation of the up-/down- coregulated gene number for the permutated domains with the same gene number. The p-values were calculated by distribution function of standard normal distribution. If the number of up-/down- coregulated genes in the domain is larger than in 95% of the background (p<0.05), the domain is defined as up-/down- coregulated. Only genes expressed at least one of the tissues are used for analysis. Fisher exact test was used to test whether there are more coregulated domains than the randomized control. Different from that of reported mammalian TADs that the coregulated domains are around 2-fold more than the random ones, our analysis indicated that the co-regulated domain are similar to the randomized control.

### Chromatin loop analysis

Chromatin loops were identified by HICCUPS from the juicer package using 10 kb bin with default parameters, only loops within 2 Mb were kept. Tissue-specific loops were identified by HICUPPSDiff with default parameters (Durand et al. 2016).

To test whether the maize loops are conserved in other species, we identified the maize loop gene pairs and their synteny genes in sorghum, rice and foxtail millets. We then calculated the physical distance of their synteny gene pairs. If a loop contains more than two genes, only the shortest distance is counted. We randomly reshuffled gene positions in the genome 1000 times cyclically and use the actual loop positions to find the randomized gene pairs as control. The control gene pairs’ synteny gene pairs in sorghum, rice and foxtail millets were identified as described above, and their physical distances are calculated. Wilcoxon rank sum test was used to examine the physical distance distribution between the actual and the randomized control gene pairs in sorghum, rice and foxtail millets.

To test whether the loop genes have higher expression correlation than randomized genes with the same physical distance but not associated with loop, we calculated the gene expression Pearson correlation of genes located in the same loop across different tissues. We randomly reshuffled gene positions in the genome 2000 times cyclically and use the actual loop position to identify the randomized loop genes as control. Their gene expression Pearson correlation in different tissues was also calculated as mentioned above. Next, we compared the two groups of distributions by Kolmogorov-Smirnov test. Same analysis was performed using the human loop genes (Rao et al. 2014).

## Acknowledgments

This work was supported by National Key Research and Development Program of China 2016YFD0101003, NSFC 91435108 and Hong Kong UGC GRF 14104515 and 14108117, Area of Excellence Scheme (AoE/M-403/16), as well as the Taishan Pandeng program. Sequencing data have been deposited in the NCBI Sequence Read Archive under the accession number PRJNA486213.

## Author contributions

S.Z. designed the research; SZ and X.T. performed the experiments; P.D., H.L., P.L. analyzed the data and; D.G., and S.Z. wrote the paper.

